# The relationship of body condition, superoxide dismutase and superoxide with sperm performance

**DOI:** 10.1101/556548

**Authors:** Christopher R. Friesen, Simon P. de Graaf, Mats Olsson

**Affiliations:** University of Wollongong, School of Earth, Atmospheric and Life Sciences, Wollongong, NSW 2522, Australia; The University of Sydney, Faculty of Science, School of Life and Environmental Sciences, NSW 2006, Australia; University of Gothenburg, Department of Biological & Environmental Sciences, Gothenburg, Sweden

**Keywords:** Sperm performance, oxidative stress, polymorphism, body condition, condition dependence, lizard

## Abstract

Sperm competition theory predicts a negative correlation between somatic investment in traits that aid in pre- and postcopulatory sexual selection. Sperm performance is critical for postcopulatory success but are susceptible to damage by free radicals such as superoxide radicals generated during mitochondrial respiration (mtSOx). Males can ameliorate damage to spermatozoa by investing in the production of antioxidants, like superoxide dismutase (SOD), which may act as a mechanistic link for pre and postcopulatory trade-offs. Some male Australian, colour-polymorphic painted dragon lizards (*Ctenophorus pictus*) possess a yellow throat patch (bib) that females prefer over non-bibbed males and are also more likely to win male-male contests indicating that males with bibs are better at monopolizing females. We tested whether the sperm performance in non-bibbed males was superior to that of bibbed males as predicted by sperm competition theory. We show that blood cell mtSOx levels are negatively correlated with SOD activity in the plasma in all males early in the breeding season but SOD was lower in bibbed males. Non-bibbed males maintain a positive correlation between body condition and SOD activity over time while bibbed males do not. Overall sperm performance was not different between the bib-morphs, however, higher mtSOx levels were negatively correlated with sperm performance in bibbed males, but not of non-bibbed males. Together these data suggest physiological associations between body condition, SOD activity and sperm performance are linked to the expression of a yellow gular patch, which may be related to intrinsic differences in metabolism of bibbed versus non-bibbed males.

**Lay summary:** Damage-inducing reactive oxygen species (ROS) are a by-product of oxygen-based energy production that are quenched by energetically expensive antioxidants. Male sexual colouration requires investment of energy and resources, which may constrain allocation to other functions like antioxidant production or spermatogenesis. Here we explored whether the body condition of colourful male lizards reflected their investment in antioxidants and reduction of ROS, which may influence sperm performance. We found that drab males in better condition had more antioxidants. Colourful males had lower levels of antioxidants and their sperm performed poorly at higher ROS levels. These results suggest a trade-off between colour maintenance and sperm performance.

## Introduction

Theoretical models of ejaculate investment under sperm competition predict an allocation trade-off between pre- and postcopulatory traits generating a negative correlation between these two types of traits (Parker, 1970; Parker et al., 2013; Parker and Pizzari, 2010; Tazzyman et al., 2009). Among individuals of the same species, nutritional resources and energy allocation—quality or condition—is especially likely to mediate pre- and postcopulatory trade-offs (Tuni et al., 2018). Indeed, empirical results that indicate investment in precopulatory traits are often negatively correlated with testes size, predominantly in species where males are able to monopolise females (Dines et al., 2015; Dunn et al., 2015; Fitzpatrick et al., 2012; Kahrl et al., 2016; Lüpold et al., 2014; Simmons and Emlen, 2006). However, in addition to size, other testicular adaptations may also be important for postcopulatory sexual selection (Ramm and Schärer, 2014).

A critical function of the testis is to protect sperm from oxidative damage during spermatogenesis and subsequent storage (Aitken and Roman, 2008; Ko et al., 2014; Ramm and Schärer, 2014). Oxidative damage of functionally critical macromolecules (membrane lipids, cellular proteins, and DNA) by reactive oxygen species (ROS) adversely affects whole organism physiological performance, health and lifespan (Finkel and Holbrook, 2000). ROS are free radicals naturally and principally produced in mitochondria during oxidation-reduction reactions occurring in the electron transport chain (ETC) during oxidative-phosphorylation generating ATP and therefore tightly linked to energy production (Murphy, 2009; Turrens, 2003). Spermatozoa are particularly vulnerable to oxidative damage because their membranes are composed of a high proportion of easily oxidized polyunsaturated fatty acids (Sanocka and Kurpisz, 2004). Therefore, selection on energetically expensive mechanisms to reduce oxidative damage is important both for life-history evolution and sexually selected traits, potentially generating condition dependent soma-germline trade-offs (Dowling and Simmons, 2009; Metcalfe and Alonso-Alvarez, 2010; Monaghan et al., 2009; von Schantz et al., 1999).

As they are terminally differentiated and transcriptionally inactive with limited antioxidant defenses, spermatozoa have limited capacity to repair molecular damage (Tremellen, 2008). Consequently, ROS production and damage are self-perpetuating in spermatozoa leading to reduced sperm function, velocity and percent of progressively motile sperm, all of which are strongly associated with male infertility in humans (Aitken et al., 1989; Aitken and Graves, 2002; Koppers et al., 2008; Tremellen, 2008). Intuitively, sperm performance is conjoined with male reproductive success so oxidative stress at the organismal level should have important implications in postcopulatory sexual selection (Firman and Simmons, 2010; Fitzpatrick and Lupold, 2014; Møller, 1988; Simmons and Fitzpatrick, 2012). Sperm competition puts males in a double bind. The production of more sperm requires increased metabolic activity in the testes (Gomendio et al., 2011; Parapanov et al., 2008; Tourmente et al., 2011), which may then be accompanied by a concomitant increase in ROS production and adaptations to ameliorate the associated costs (delBarco-Trillo and Roldan, 2014; Ribou and Reinhardt, 2012). Even in species with relatively short-term sperm storage and with limited opportunity for sperm competition, such as humans (Simmons et al., 2004; van der Horst and Maree, 2014), oxidative damage strongly influences sperm performance and male reproductive success (Aitken et al., 2014). Allocating resources for protecting sperm from oxidative damage to maintain high sperm performance is even more vital in the context of sperm competition (Dowling and Simmons, 2009).

Protection from and repairing damage caused by ROS during spermatogenesis is potentially costly and may be condition dependent (Costantini, 2008, 2014; Monaghan and Costantini, 2014). There is increasing evidence of the conditional dependence of sperm production (e.g. numbers and morphology Dávila and Aron, 2017; Kahrl and Cox, 2015) and sperm velocity, motility and fertilization success (Mitre et al., 2004; Rakitin et al., 1999). Male condition dependence of oxidative status affects sperm quality, and there is increasing evidence that male colouration reflects antioxidant status (Helfenstein et al., 2010; reviewed in, Svensson and Wong, 2011; von Schantz et al., 1999) and studies indicate colouration and sperm quality can be correlated (Locatello et al., 2006; Peters et al., 2004; Pitcher et al., 2007; Rowe et al., 2010). For example, Helfenstein et al. (2010) showed that male great tits (*Parus major*) with paler carotenoid-based breast plumage subjected to experimentally elevated workloads had higher lipid peroxidation (an indicator of oxidative damage) as well as reduced sperm motility and ‘swimming ability’; these effects were ameliorated with supplementation of dietary carotenoid antioxidants. Age-related declines in the innate antioxidant capacity of red junglefowl (*Gallus gallus*), was associated with an increase in oxidative damage, and a decline in sperm quality (Noguera et al., 2012). Decreased endogenous antioxidant expression translates to lower sperm competitiveness as Garratt et al. (2013) show that mice deficient in the endogenous antioxidate superoxide dismutase (SOD) had no fertilization success in sperm competition trials. Thus, the role of oxidative stress in sperm performance and/or fertilization success *in vivo*, preferably in wild populations, merits attention (Dowling and Simmons, 2009). Such analyses would be particularly interesting in intraspecific studies on males with divergent reproductive strategies characterized by trade-offs between pre- and postcopulatory sexually selected traits (Lüpold et al., 2014).

Colour-polymorphic species can be a useful tool in determining the selective forces that drive sexually selected trait expression in wild populations. Heritable colour morphs often have divergent behaviours and physiology, with associated differences in reproductive tactics in animals with an otherwise common genetic background (Healey et al., 2007; Huxley, 1955; Olsson et al., 2007b; Olsson et al., 2009a; Pryke et al., 2007; Pryke and Griffith, 2006; Sinervo and Lively, 1996; reviewed in Wellenreuther et al., 2014).

Here we assess the relationship between an important mitochondrial ROS—antioxidant pair, a measure of resources (body condition: BCI) and sperm performance in a wild colour-polymorphic lizard, the Australian painted dragon (*Ctenophorus pictus*). Primary ROS production in mitochondria usually begins with the single reduction of molecular oxygen (O_2_) to form superoxide (O^•-^), which is the rate limiting step in the subsequent damaging cascade of ROS production (Andreyev et al., 2015; Turrens, 2003). On its own, superoxide is a relatively weak oxidizer of macromolecules; however, superoxide reduces iron (Fe^2+^) which may then catalyse the disproportionation of hydrogen peroxide (H_2_O_2_) to produce the hydroxyl radical (^•^OH), which is the strongest oxidizing agent in biological systems (Fridovich, 1995; Turrens, 2003). Superoxide that is not quenched readily exits cell-membranes with high concentrations of anion channels, such as erythrocytes (Fridovich, 1995; Lynch and Fridovich, 1978). So we will focus on superoxide and its elimination by the action of an endogenous enzyme, superoxide dismutase (SOD), which, at sufficiently high concentrations, eradicates O^.-^ at the rate of diffusion (Fridovich, 1995; Turrens, 2003). Thus, although there are other important endogenous antioxidants (e.g., catalase and various isoforms of glutathione peroxidase), the balance between superoxide and SOD is a significant mediator of incipient oxidative stress during energy (ATP) production.

Oxidative stress and sexual selection biology of painted dragons have been the subject of extensive investigations. For example, aggressive colour-morphs have higher testosterone (T), and superoxide levels than less aggressive males that tend to have lower T (Olsson et al., 2007a; Olsson et al., 2009c). Although colour is in part carotenoid-based, a potential exogenous antioxidant, dietary manipulation of carotenoids does not mediate a relationship between superoxide and bright colouration (Olsson et al., 2008a). However, experimental reduction of superoxide levels using exogenous doses of the SOD mimetic, EUK 134, does reduce colour fading over the breeding season (Olsson et al., 2012b). Males in better body condition (residuals of mass-length regression) have enhanced morph-specific colour maintenance despite higher net ROS levels (Friesen et al., 2017b) indicating that investment in endogenous antioxidants, such as SOD, may be condition-dependent. Furthermore, baseline superoxide level varies among individuals and is heritable (Olsson et al., 2008b), but whether this is due to reduced production of ROS or increase quenching of ROS by endogenous antioxidants in unknown. Collectively these studies suggest that there are evolutionarily important links between body condition, SOD and superoxide levels, and sexual colouration in painted dragons.

In this paper we exploit natural variation in a colour trait, the presence or absence of a yellow gular patch (henceforth, bib **Figure 1**), which is an important, but costly, sexually selected colour-trait in painted dragons. Bibbed male painted dragons experience greater declines in body condition in the wild than non-bibbed males (Healey and Olsson, 2009a; Olsson et al., 2009a). Males with bibs also have greater telomere attrition (Rollings et al., 2017) and higher resting metabolic rate through the breeding season than non-bibbed males (Friesen et al., 2017a). These costs of bearing a bib, must come with potential fitness benefits to explain the trait’s persistence in the population. Indeed, bibbed males are more attractive to females in staged preference trials and more likely to win male-male competition trials when pitted against non-bibbed rivals indicating there is a precopulatory mating advantage for bibbed males (McDiarmid et al., 2017b). Bibbed males are less likely than non-bibbed males to lose paternity to neighbouring males in the wild (Healey and Olsson, 2009a; Olsson et al., 2009a), which may be due to efficient mate monopolization. McDiarmid et al. (2017b), hypothesized that this precopulatory advantage of bibbed males would relax postcopulatory selection on sperm performance, however sperm motility and velocity did not differ between bibbed and non-bibbed males. McDiarmid et al. (2017b) did not measure superoxide or SOD. We make three straightforward predictions: 1) SOD and superoxide level should be negatively correlated, 2) sperm performance and superoxide levels should be negatively correlated, and 3) if SOD is indeed costly, body condition should be positively correlated with SOD levels. Finally, if bibbed males face a resource allocation trade-off, then bibbed males should have lower sperm performance after accounting for SOD and body condition than non-bibbed males.

**Figure 1.**
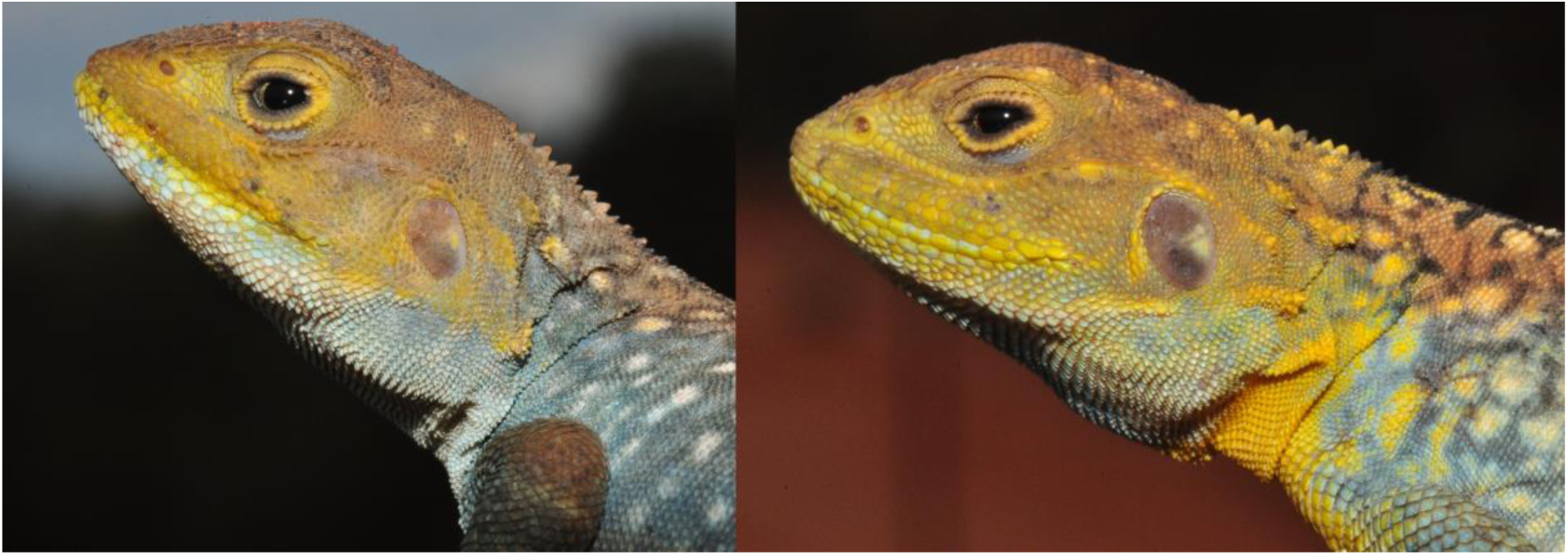
Male painted dragon lizards. On the left is a male without a yellow throat patch (i.e. a non-bibbed male) and on the right is a male with a yellow throat patch (i.e. a bibbed male).

## Methods

### Animal collection/housing and mating trials

The Australian painted dragon (*Ctenophorus pictus*, Peters 1866) is a small (adult: mass 8 – 18 g; snout-to-vent length [SVL] 55-75mm) agamid lizard native to arid parts of central New South Wales to Western Australia (Cogger, 2014). All research was conducted with the approval of the University of Sydney Animal Ethics Committee Project Number: 2013-2016/6050 and were collected under NSW National Parks and Wildlife Service Scientific licence number SL100352. Lizards were caught at Yathong Nature Reserve, in western New South Wales (145°C35′; 32°C35′) and were brought back to holding facilities at the University of Sydney (14, October: N = 40 females; N = 56 males; 20 bibbed and 36 non-bibbed males). The animals were sampled at random at the study site and, hence, approximately represent natural frequencies in the wild at the time. Males were kept individually in tanks (60 × 60 × 50cm) on a 12:12 h (light: dark) light regime with a basking light for thermoregulation. They were watered daily, and fed crickets and mealworms to satiation every second day.

After a three-week acclimation period, we conducted mating trials on five different days in the period between 6-Nov.-4 Dec (5-8 matings per day). This period corresponds with the middle third of the breeding season in the wild (late September-late December). We obtained 5-8 matings on any given day which allowed for timely processing of both sperm and blood samples with minimal temporal separation between them. Each male was observed continuously after the introduction of a receptive female to his own cage. Males usually mated within 10 minutes of the female’s introduction, and trials were terminated if mating did not occur after 1 hour. Only those males that mated were included in this study (N = 31 of 56 males mated; 12 bibbed and 19 non-bibbed males). Although not all males mated, the ratio of bibbed/non-bibbed males was the same as those brought back to the lab (X^2^_df 1_ = 0.077, P = 0.781). Females were used only once during the receptive phase of their 3 week reproductive cycle (Olsson et al., 2009b).

Subsequent to copulation, semen was collected from the female’s cloaca by pulling the ejaculate into a 1 mL syringe (Olsson, 2001) preloaded with 100 µL Hams F-10 medium (Cat # 99175, Irvine Scientific, Santa Ana, CA, USA; 21 mM HEPES buffer, 4 mM sodium bicarbonate, 1 mM calcium lactate, 0.5 mM magnesium sulfate, 5 mg mL-1 (0.5%) human albumin; e.g., (Friesen et al., 2013; Friesen et al., 2014; Mattson et al., 2007; McDiarmid et al., 2017b). Sperm samples were kept at 35±1 °C in an incubator until they were analysed using computer-assisted sperm analysis (CASA, see below) ≤ 30 minutes after collection; 35 °C is the preferred body temperature of this species (Melville and Schulte II, 2001). A pilot study (N = 6) showed no significant decline in sperm motility or velocity after 60 minutes, and was verified in the full data set presented in this paper (see results below). We collected a blood sample in a capillary tube from a vessel in the corner of the mouth, the *vena angularis*, after it was gently punctured with the tip of an 18 ga syringe needle. Each blood sample was taken within 5 min of semen collection for mitochondria superoxide and plasma superoxide dismutase analysis (below). Body mass and SVL were recorded the same day as mating trials.

### Sperm performance—Computer aided sperm analysis (CASA)

The ejaculate was diluted to 1×10^6^ / mL with Hams F-10 medium and slowly pipetted (10.0 μL) into the chamber of a pre-warmed sperm analysis slide (Hamilton-Thorne 2X-CEL^®^; Beverly, MA, USA; warmed to 35 °C on a slide warmer). Sperm were examined using CASA at a frame rate set to 60 hz (Hamilton-Thorne, IVOS-II; Beverly, MA, USA). Sperm motility was determined for a minimum of 300 sperm per individual. In pilot work, we found it impossible to eliminate all particulates from the ejaculate without negatively affecting sperm motility with excessive washing. Therefore, we modified CASA factory settings to classify sperm as motile only if its velocity was ≥5 µm/s with no constraints on path straightness. In addition, we visually inspected each sperm track to verify that only sperm were included in the analysis and to confirm accurate tracking of multiple sperm with intersecting paths at the time of analysis. In addition to providing proportion of motile sperm (MOT), we used the following standard average measures of sperm kinematics data generated by CASA for each sample for PCA analysis (see “statistics” below): straight-line velocity (VSL), curvilinear velocity (VCL), average path velocity (VAP), amplitude of lateral head displacement (ALH), beat cross frequency (BCF), linearity of movement (LIN), and proportion of progressively motile sperm (PMOT). Progressively motile sperm (i.e., sperm that were ‘going somewhere’ and not merely convulsing or turning in place) were defined as sperm with an average path velocity (VAP) of ≥50µm/s and a straightness index (STR) of 70% (VSL/VAP ≥70%).

### Mitochondrial Superoxide—Flow-cytometry fluorescence-activated cell sorting (FACS)

We followed previously published protocols for measuring mitochondrial superoxide (Ballen et al., 2012; Friesen et al., 2017b; Giraudeau et al., 2016). Briefly, an aliquot of a single sample of whole peripheral blood (10 μl) was diluted immediately with 9 volumes of phosphate buffered saline (PBS; 137 mM NaCl, 2.7 mM KCl, 1.5 mM KH2PO4, 8 mM Na2HPO4, pH 7.4) and stored on ice prior to FACS analyses that were completed within 2 hours of sampling. The remainder of the blood sample was kept on ice until centrifugation for plasma collection and subsequence storage at −80°C (3 hours after blood sampling) for later quantification of SOD. Prior to staining blood cells for mt-superoxide, the diluted blood was further diluted 50-fold in PBS and then centrifuged (300 g for 5 min) to pellet blood cells. Cells were resuspended in 100 μl of PBS containing 5 μM MitoSOX Red (MR; Molecular Probes, Invitrogen, Life Technologies, Sydney, Australia). MR was added from stock solutions in dimethyl sulfoxide (DMSO); the final concentration of DMSO was 0.2% (v/v) or less. Cells were subsequently incubated at 35 °C for 30 min then washed with PBS by centrifugation as described above and held on ice until analysed by flow cytometry; 50,000 events (cells) were acquired for all samples. Flow cytometry was performed using an Accuri C6 flow cytometer (BD Accuri Cytometers, Ann Arbor, MI, USA) with excitation at 488 nm and emitted fluorescence collected using band pass filters of 575 ± 13 nm. Data were acquired and analysed using BD Accuri C6 Software. On the basis of forward angle laser scatter and side angle laser scatter, a number of blood cell populations were discerned; the results obtained were similar for all these populations. For each sample, the arithmetic mean fluorescence for all 50,000 cells acquired was determined using Accuri C6 software and used to compare between samples and treatments. (A pilot experiment determined that the particulate matter in the semen samples precluded measurement of mitochondrial superoxide production in sperm with flow cytometry).

### Superoxide dismutase activity

We followed the same protocol described previously in this species (Olsson et al., 2012a; Olsson et al., 2012b) that assayed SOD in blood plasma samples collected within five minutes of semen collection. Briefly, we used the Superoxide Dismutase Assay Kit II (catalogue no. 574601, Calbiochem, supplied by Merck Pty Ltd., Victoria, Australia). This kit is designed to generate superoxide radicals with xanthine oxidase and hypoxanthine. The superoxide radicals are dismutated by SOD in the plasma sample to molecular oxygen and hydrogen peroxide and detected by tetrazolium salt. This SOD assay measures all three types of SOD (Cu/Zn-, Mn- and Fe-SOD). Absorbance measurements were taken using a microplate reader (Pherastar^®^FSX, BMG Labtech, Mornington, Victoria, Australia). The optimal plasma sample dilution was determined to be a factor of 48 in previous studies (Olsson et al., 2012b), indicating that SOD is abundant in the plasma of this species. Standards and samples were run in duplicate and were placed randomly on the plates for intra-assay precision. The intraclass correlation coefficient (ICC) between two samples from the same individual was ICC = 0.880, P < 0.001. One unit of SOD activity is defined as the amount of enzyme needed to exhibit 50 % dismutation of the superoxide radical to hydrogen peroxide.

### Statistical analysis

#### Data preparation

We calculated body condition (BCI) as the standardized residuals (mean = 0; standard deviation =1) from linear regressions of ln(body mass) as a function of ln(SVL) in all males, not just those that mated (Olsson et al., 2009a).

After inspection of normal Q-Q plots and the failure of Shaprio-Wilk tests of normality, mean fluorescence values of mitochondrial superoxide (henceforth “mtSOx”: W = 0.858, P < 0.001) and superoxide dismutase activity (SOD: W = 0.883, P = 0.003) were ln-transformed to improve normality (ln(mtSOx): W = 0.961, P = 0.317; ln(SOD): W = 0.993, P = 0.999) and reduce heteroscedasticity. Hereafter, mtSOx and SOD refer to these transformed variables unless otherwise stated.

A priori, we wanted to assess proportions of motile (MOT) and progressively motile (PMOT) sperm. MOT is one of the most commonly reported sperm performance parameters while PMOT is a useful holistic indicator of efficient mitochondrial function, membrane integrity and fertilization success in most species studied to date, including for example, humans, horses, ram, rabbit and poultry (De Graaf et al., 2006; Froman and Feltmann, 2000; Froman and Kirby, 2005; Johinke et al., 2014; Quintero-Moreno et al., 2003; Rickard et al., 2014; Simon and Lewis, 2011). PMOT captures both velocity and path straightness and also scales to the relative population size of progressively motile sperm in the ejaculate. However, it is unclear how these two variables and the other six kinematic parameters generated by CASA are related.

Initial data exploration with Q-Q plots indicated MOT and PMOT were normally distributed, which was also confirmed using Shaprio-Wilk tests of normality (MOT: W = 0.946, P = 0.123; PMOT: W = 0.976, P = 0.684). Furthermore, MOT, VSL, VCL, VAP, ALH, BCF, LIN, and PMOT were correlated (|r| range: 0.276-0.973). Rather than run multiple separate analyses with each sperm kinematics parameter, following Helfenstein et al. (2010), we chose to reduce these eight dimensions into one variable using principle component analysis with varimax orthogonal rotation. The first axis (PC1), which we call “sperm performance” and Helfenstein et al. (2010a) call “swimming ability”, explained 69.73% of the variance in the kinematic parameters. All kinematic parameters were significantly correlated with PC1 after rotation (Listed here in order of increasing strength of positive correlation with PC1: LIN, ALH, MOT, PMOT, VCL, VAP, VSL; r ranged 0.663-0.994, all p <0.0001; BCF (beat frequency) had a significantly negative correlation with PC1 r = −0.578, P< 0.0001). Thus, individuals with relatively large and positive values of PC1 can be described as having proportionally more sperm that swim relatively fast and straight and were more efficient given the negative correlation with beat frequency.

#### Significance tests

We report basic comparisons of mass, SVL and body condition between bib-morphs at time of capture and at mating. We pre-planned to fit separate models including one covariate of interest (BCI, mtSOx, SOD) and its interaction with bib-morph. Therefore, we chose to conduct significance tests on three separate relationships using restricted maximum likelihood (REML) estimated Generalized Linear Models (GLM; normal distribution, identity link function) with Satterthwaite approximated degrees of freedom and robust covariance estimation to manage unbalanced sample sizes in IBM^®^ SPSS^®^ version 24. We report Likelihood ratios and associated p-values from Omnibus tests of full versus intercept-only models for completeness but base our interpretations on parameter estimates for the pre-planned comparisons (Rutherford, 2011). All full models reported here initially included main effects of bib-morph and the covariate of interest as well as their interaction, but the interaction parameters were dropped if estimates were not significant at α > 0.05.

#### Figures

We included separate regression lines for each ‘morph’ in all plots regardless of significance. All plots and R^2^ values (when reported) for these simple linear regressions (SLR) were generated in SigmaPlot^®^ 13.0 (Systat Software, Inc. San Jose CA, USA).

## Results

Of the males that mated and were used for sperm collection, bibbed males were slightly but significantly longer (SVL: 66 mm ± 0.07 se) than non-bibbed males (SVL: 68 mm ± 0.07 se) at capture: F_1, 29_ = 5.578, P = 0.025); SVL did not change in the acclimation period. Bibbed and non-bibbed males did not differ significantly in mass or body condition at the time of capture or mating (all P ≥ 0.119) although all males gained an average of 0.626 g ± 0.229 se or 4.7% increase in body mass in this period (F_1,29_ = 6.820, P = 0.014).

### Relationship between body condition and superoxide dismutase activity

Superoxide dismutase activity, measured at the time of mating, was positively related to body condition at time of capture (SOD~ BIC_capture_: r = 0.336, F_1, 29_ = 4.465, P = 0.043). Although BCI increased from time of capture to time of mating in the lab, SOD activity decreased slightly over the course of the mating trials (i.e., sampling period) for all males (SOD ~ date: r = −0.355, F_1,29_ = 4.193, P = 0.050) regardless of bib phenotype (date × Bib, P = 0.748). At the time of each individual male’s mating, BCI was not significantly correlated with SOD activity, however, SOD activity was positively related to BCI in non-bibbed males, but not in bibbed males (Bib × BCI_mating_, P =0.015; **Figure 2b**,**Table 1**), and non-bibbed males had higher SOD activity than bibbed males (P = 0.041).

**Table 1:** Model effects of GLM (normal, identity link function) SOD activity as a function of bib and BCI at the time of mating. Full model: ln(SOD) ~ Bib + BCI_mating_ + Bib * BCI. Likelihood ratio χ^2^_df3_ = 8.9166, P = 0.043 vs intercept only model. Bib (n = 12), Non-bibbed (n = 19), a positive coefficient indicates non-bibbed greater than bibbed. Bold text indicates P ≤ 0.05.

| $\ln(\text{SOD}) \sim$ | $\beta(\text{se})$ | 95% Likelihood CI | Type III Sum of Squares |
| --- | --- | --- | --- |

| Source |  | (lower, upper) | Wald | Sig. |
| --- | --- | --- | --- | --- |
| Intercept | -1.951(0.165) | -2.243, -1.658 | 328.891 | **<0.001** |
| Bib | 0.394(0.194) | 0.021, 0.768 | 4.161 | **0.041** |
| BCI<sub>mating</sub> | -0.161(0.129) | -0.413, 0.90 | 1.570 | 0.210 |
| Bib x BCI<sub>mating</sub> | 0.407(0.168) | 0.028, 0.786 | 5.878 | **0.015** |

**Figure 2.**
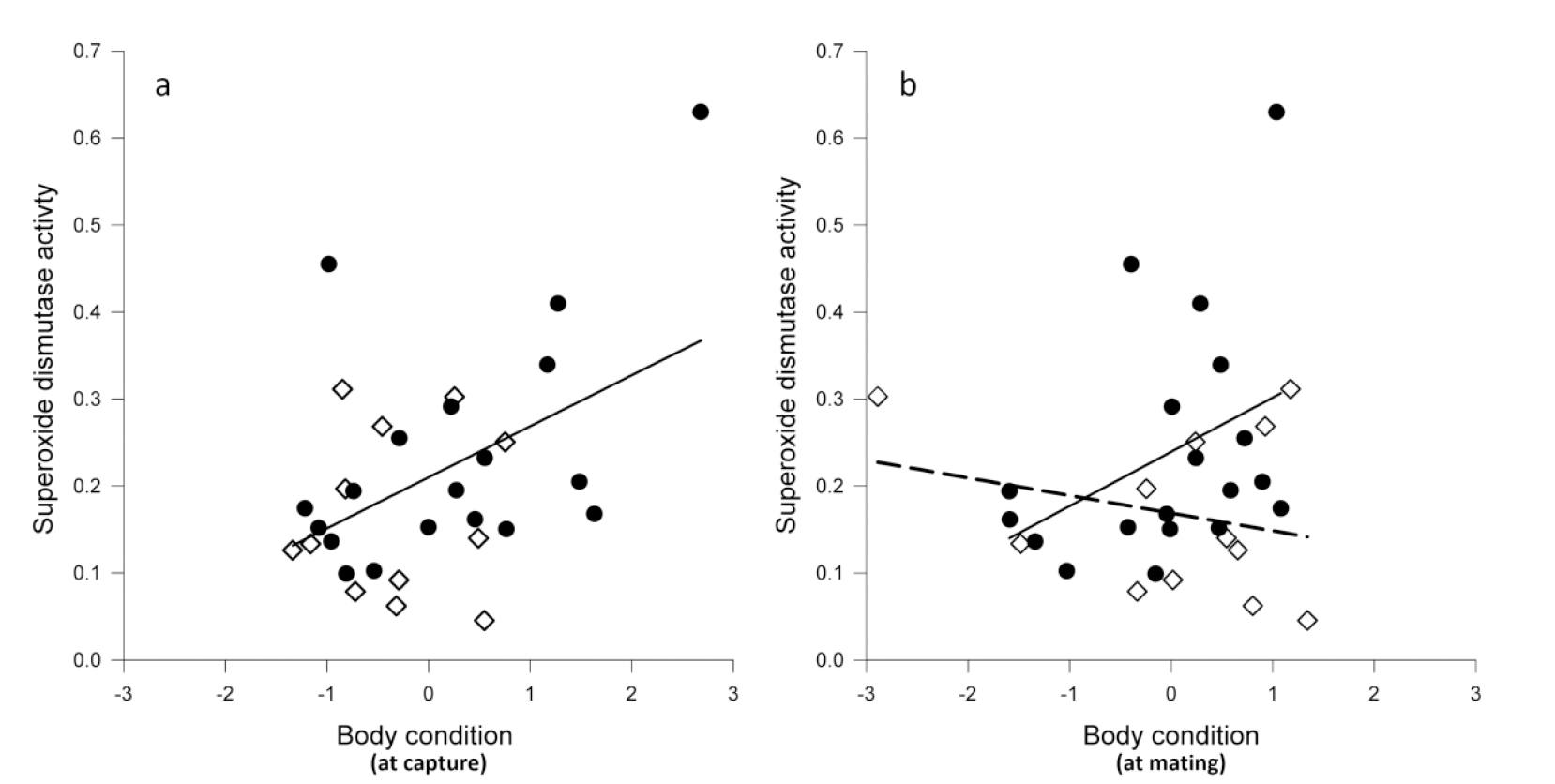
Relationship between SOD activity (untransformed data graphed) and body condition (BCI, residual ln(body mass) given ln(snout to vent length)). In each graph, solid circles (•) represent non-bibbed males and unfilled diamonds (◊) represent bibbed males. Panel **a)** SOD activity at mating as a function of body condition *at capture*: R^2^ = 0.213, P = 0.009. Panel **b)** SOD activity at mating as a function of body condition at mating: separate regression lines for each morph are plotted due to the significant morph × BCI interaction (P = 0.015; solid line represents non-bibbed males (R^2^ = 0.149, P = 0.103), dashed line represents bibbed males (R^2^ = 0.066, P = 0.420).

### Relationship between mitochondrial superoxide (mtSOx) and SOD

Overall, mtSOx tended to decrease with increasing SOD activity (P = 0.012) **Table 2; Figure 3**. Superoxide levels increased significantly over the course of the sampling period in all males (r = 0.564, F_1, 29_ = 13.526, P = 0.001) regardless of bib phenotype (date × Bib, P = 0.371).

**Table 2:** Model effects of GLM (normal distribution, identity link function) mtSOx levels as a function of bib and SOD activity at the time of mating. Full model: ln(mtSOx) ~ Bib + ln(SOD) + Bib * ln(SOD): Likelihood ratio χ^2^ _df3_ = 6.345, P = 0.096 vs intercept only model. Reduced model: mtSOx~ln(SOD) Likelihood ratio χ^2^_df1_ = 4.646, P = 0.031 Bib (n = 12), Non-bibbed (n = 19), a positive coefficient indicates non-bibbed greater than bibbed. Bold text indicates P ≤ 0.05.

| Ln(mtSOx)~<br>Source | $\beta(\text{se})$ | 95% Likelihood CI<br>(lower, upper) | Type III Sum of Squares | |
| --- | --- | --- | --- | --- |
|  |  |  | Wald | Sig. |
| Intercept | 6.786(0.470) | 5.646, 7.926 | 445.140 | **<0.001** |
| Bib | 1.174(0.699) | 0.294, 2.642 | 2.820 | 0.093 |
| $\ln(\text{SOD})$ | -0.701(0.194) | -1.256, -0.147 | 6.297 | **0.012** |
| Bib x $\ln(\text{SOD})$ | 0.527(0.349) | -0.265, 1.318 | 2.276 | 0.131 |

**Figure 3.**
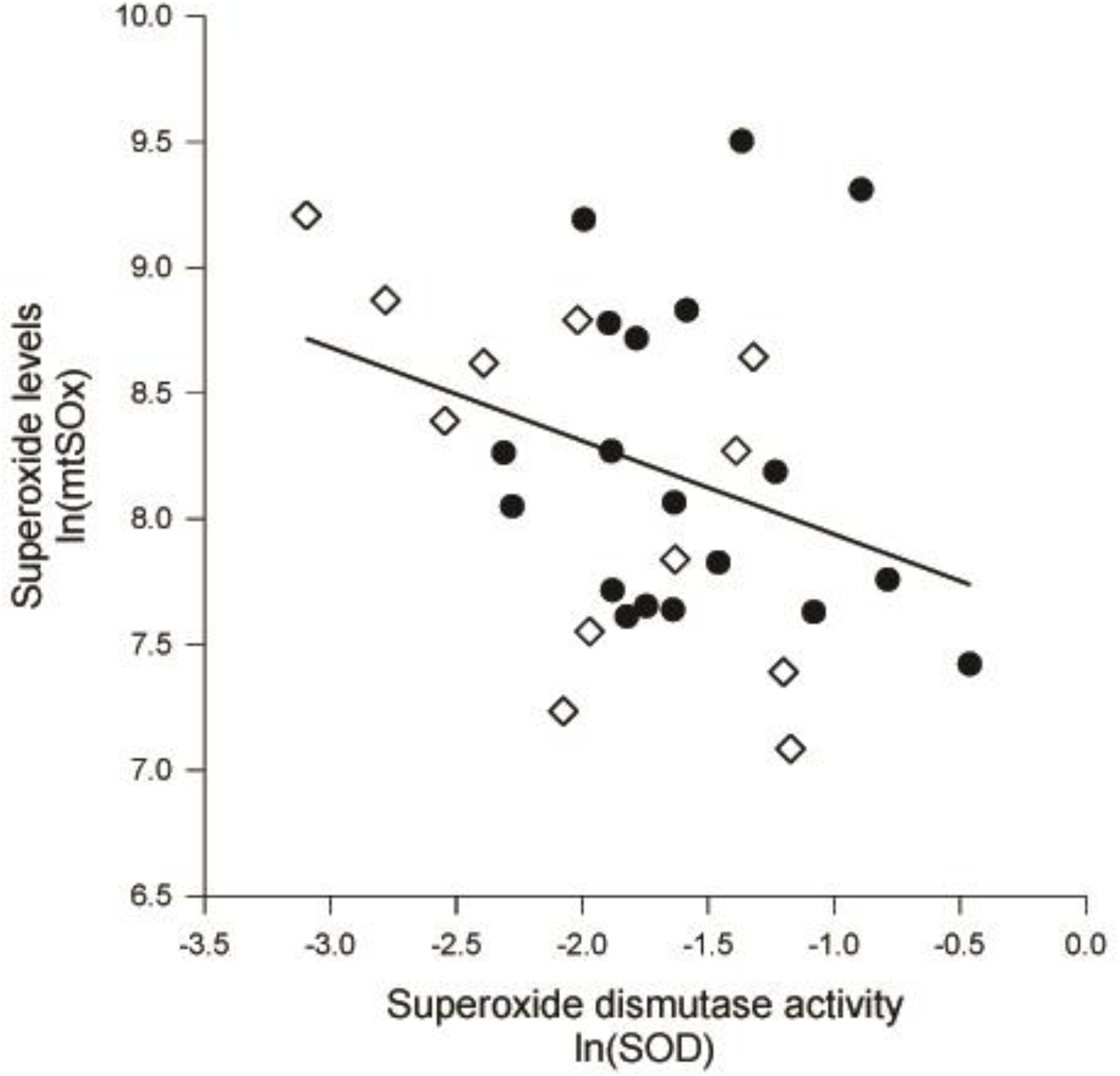
Negative relationship between mitochondrial superoxide levels in blood cells and plasma superoxide dismutase activity (R^2^ = 0.107, P = 0.018). Solid circles (•) represent non-bibbed males and unfilled diamonds (◊) represent bibbed males.

### Relationship between mitochondrial superoxide and sperm kinematic parameters

We constructed GLMs with bib, superoxide levels and their interaction as independent variables and three different kinematic parameters as response variables: MOT, PMOT and PC1 or “sperm performance” **Tables 3a-c** respectively. Sperm motility was significantly and negatively related to superoxide levels (P = 0.001), but motility did not differ between bib-morphs (P = 0.358) **Figure 4**. Sperm performance (PC1) and the proportion of progressively motile sperm (PMOT) was negatively linked with superoxide levels (mtSOx: P ≤ 0.004). Progressive motility of sperm declined in both morphs, but significantly more steeply in bibbed males (bib × mtSOx P = 0.036: **Figure 5a**). While sperm performance (PC1) declined with increasing mtSOx levels in bibbed males (bib × mtSOx, P = 0.036 **Figure 5b**), no such decline occurred in non-bibbed males.

**Table 3a:** Model effects of GLM (normal distribution, identity link function) sperm motility (MOT) as a function of bib and mtSOx level at the time of mating. Full model: MOT ~ mtSOx + bib + bib*mtSOx: Likelihood ratio χ^2^_df3_ = 5.486, P = 0.139 vs intercept only model. Reduced model: MOT ~ mtSOX Likelihood ratio χ^2^_df1_ = 4.747, P = 0.029. Bib (n = 12), Non-bibbed (n = 19), a positive coefficient indicates non-bibbed greater than bibbed. Bold text indicates P ≤ 0.05.

| MOT~<br>Source | $\beta(\text{se})$ | 95% Likelihood CI<br>(lower, upper) | Type III Sum of Squares | |
| --- | --- | --- | --- | --- |
|  |  |  | Wald | Sig. |
| Intercept | 220.008(56.382) | 48.628, 391.39 | 21.279 | **<0.001** |
| Bib | -73.048(79.553) | -300.85, 154.754 | 0.843 | 0.358 |
| mtSOx | -19.993(6.822) | -40.919, -0932 | 11.244 | **0.001** |
| Bib x mtSOx | -8.231(9.470) | -19.485, 35.948 | 0.755 | 0.385 |

**Table 3b:** Model effects of GLM (normal distribution, identity link function) progressively motile sperm (PMOT) as a function of bib and mtSOx level at the time of mating. Full model: PMOT ~ mtSOx + bib + bib*mtSOx: Likelihood ratio χ^2^_df3_ = 13.218, P = 0.004 vs intercept only model. Bib (n = 12), Non-bibbed (n = 19), a positive coefficient indicates non-bibbed greater than bibbed. Bold text indicates P ≤ 0.05.

| PMOT~<br>Source | $\beta$(se) | 95% Likelihood CI<br>(lower, upper) | Type III Sum of Squares | |
| --- | --- | --- | --- | --- |
|  |  |  | Wald | Sig. |
| Intercept | 1.872(0.448) | 0.993, 2.750 | 17.445 | **< 0.001** |
| Bib | -1.015(0.502) | -1.998, -0.32 | 4.094 | **0.043** |
| mtSOx | -0.186(0.053) | -0.291, -0.082 | 12.282 | **< 0.001** |
| Bib x mtSOx | 0.013(0.003) | 0.008, 0.242 | 4.398 | **0.036** |

**Table 3c:** Model effects of GLM (normal distribution, identity link function) “sperm performance” (PC1) as a function of bib and mtSOx level at the time of mating. Full model: PC1 ~ mtSOx + bib + bib*mtSOx: Likelihood ratio χ^2^_df3_ = 6.283, P = 0.099 vs intercept only model (we did not reduce model given significant parameter estimates). Bib (n = 12), Non-bibbed (n = 19), a positive coefficient indicates non-bibbed greater than bibbed. Bold text indicates P ≤ 0.05.

| Sperm<br>performance<br>(PC1)<br>Source | $\beta$(se) | 95% Likelihood CI<br>(lower, upper) | Type III Sum of Squares | |
| --- | --- | --- | --- | --- |
|  |  |  | Wald | Sig. |
| Intercept | 20.145(7.202) | 6.030, 34.259 | 7.825 | **0.005** |
| Bib | -17.744(8.617) | -34.633, -0.854 | 4.240 | **0.039** |
| mtSOx | -2.456(0.863) | -4.147, -0.765 | 8.105 | **0.004** |
| Bib x mtSOx | 2.157(1.029) | 0.141, 4.173 | 4.398 | **0.036** |

**Figure 4.**
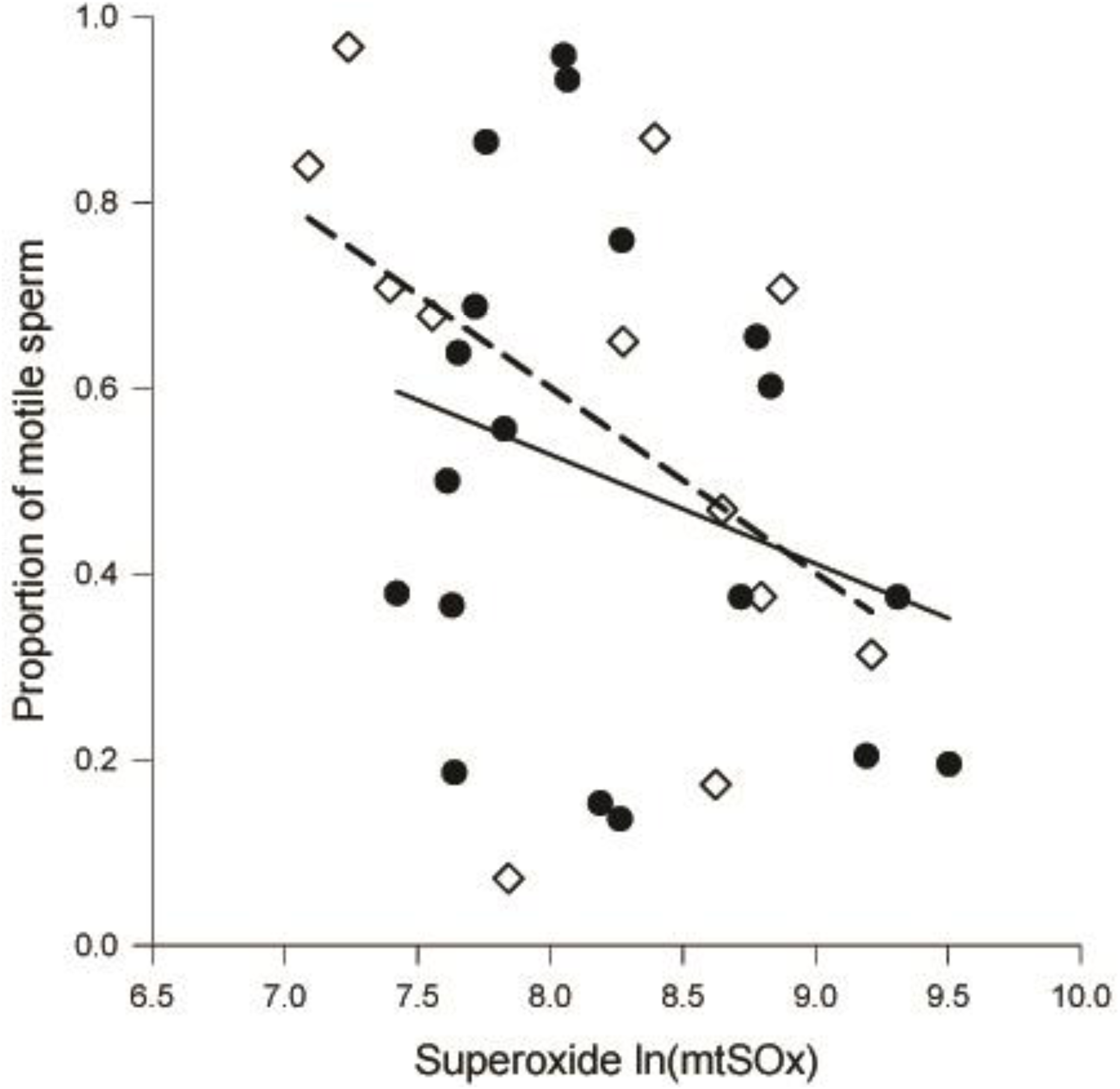
Overall negative relationship between sperm motility and mitochondrial superoxide levels (P = 0.001). We have graphed separate regressions for each bib-morph: Solid circles (•) represent non-bibbed males (solid regression line, R^2^ =0.080) and unfilled diamonds (◊) represent bibbed males (dashed regression line, R^2^ = 0.249).

**Figure 5.**
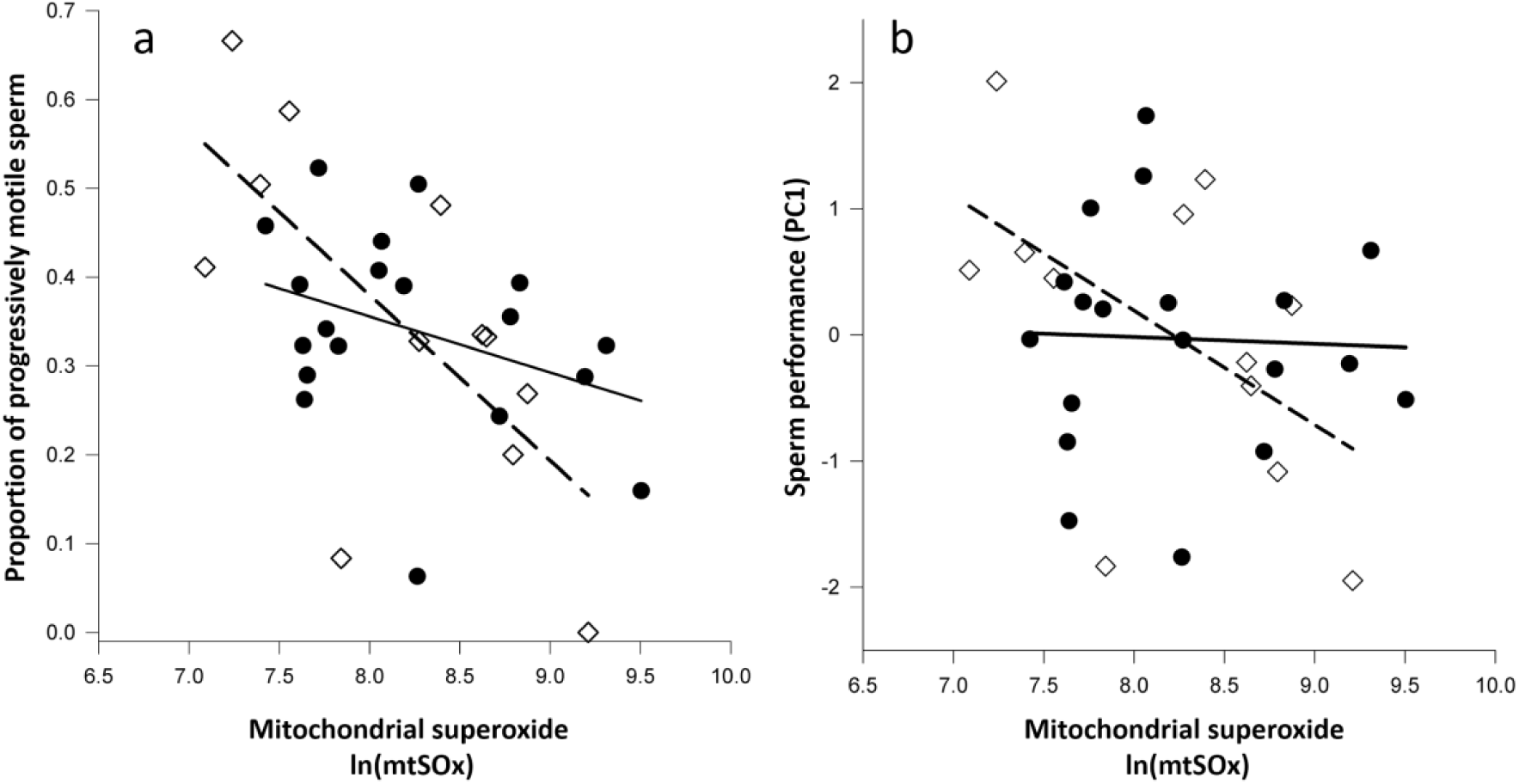
a) Negative relationship between the proportion of progressively motile sperm (PMOT) and mitochondrial superoxide levels in blood cells (ln(mtSOx); P < 0.001). Regression lines were calculated and plotted separately for each morph given the significant bib × ln(mtSOx) level interaction (P = 0.036). Solid circles (•) represent non-bibbed males (solid regression line: R^2^ = 0.130) and unfilled diamonds (◊) represent bibbed males (dashed regression line: R^2^ = 0.458). b) Negative relationship between sperm performance (PC1) and mitochondrial superoxide levels (ln(mtSOx), P = 0.019). PC1 is positively correlated with the following sperm kinematic parameters: LIN, ALH, MOT, PMOT, VCL, VAP, VSL see text for definitions (r ranged 0.663-0.994, all p <0.0001); BCF (beat frequency) had a significantly negative correlation with PC1 (r = −0.578, P< 0.0001). Regression lines were calculated and plotted separately for each morph given the significant bib × ln(mtSOx) level interaction (P = 0.038). Solid circles (•) represent non-bibbed males (solid regression line: R^2^ = 0.002) and unfilled diamonds (◊) represent bibbed males (dashed regression line: R^2^ = 0.288).

## Discussion

We investigated prospective links between body condition, antioxidant defences (SOD), whether antioxidant levels were associated with ROS levels (mtSOx) and finally whether the relationship between ROS levels and sperm performance differ between bib- and non-bibbed male painted dragon lizards. Our results suggest that somatic levels of ROS production and antioxidant protection are associated with sperm performance. Mitochondrial superoxide (mtSOx), measured in blood cells (somatic), is negatively associated with the proportion of motile sperm (MOT) and the proportion of progressively motile sperm (PMOT) in both bib-morphs (**Figures 4 and 5a**). Similar to McDiarmid et al. (2017b), we did not find bib-morph specific differences in sperm kinematic parameters per se, until superoxide was included as a covariate. There were morph-specific differences in the association of blood cell superoxide level with progressive sperm motility. The negative relationship between superoxide levels and sperm performance was most pronounced in bibbed males (represented by PMOT and PC1, **Figures 5a & 5b**). Sperm performance seems to be strongly, albeit indirectly linked to oxidative status in bibbed males, but less tightly so in non-bibbed males (**Figure 5b**). This may reflect differences in superoxide dismutase (SOD) levels in plasma (i.e., somatic investment), which were negatively related with mtSOx in both morphs, but significantly higher in non-bibbed males. Given the susceptibility of spermatozoa to ROS damage it is not surprising that there are many studies that demonstrate negative correlations between antioxidants within the semen and sperm performance in humans (Aitken et al., 1989; Aitken and Graves, 2002; Aitken and Roman, 2008; Sanocka and Kurpisz, 2004) (e.g., reviewed in Aitken, 2016); and a few in non-mammalian models (e.g., Rojas Mora et al., 2016). Nevertheless, a holistic idea of organismal function in which levels of somatic antioxidants reflects overall condition or health is also supported in the literature, as reproductive senescence and many disease states in humans are also associated with oxidative stress in somatic tissues and declines in sperm performance and infertility (Aitken, 2016; Aitken et al., 2014). The link between SOD activity and body condition we found tentatively suggests a trade-off between the allocation of resources to aggressiveness and colour investment in the soma of bibbed males on one hand, and the buffering against the negative effect of superoxide on sperm performance in non-bibbed males on the other.

As predicted, SOD activity, in both bibbed and non-bibbed males, was positively related to body condition (BCI) at the time of capture. SOD activity measured nearly a month after body condition measures were taken had a positive relationship indicating that SOD activity may depend more on the available resources at the time of SOD production, than at the time of measurement; we have recently documented similar time-lagging relationships between DNA-damage repair and telomere attrition in this species (Olsson et al., 2018). However, the relationship of SOD activity and BCI measured at the time of mating is more complex. In non-bibbed males, BCI at mating was still positively related to SOD activity, but not so in bibbed males. Indeed, SOD levels were significantly lower in bibbed males than non-bibbed males even after accounting for body condition at the time of mating.

Positive links between colour traits, oxidative stress and sperm performance have been demonstrated in a few species of birds, such as: Helfenstein et al. (2010) work on great tits mentioned in the introduction; the positive association between colour, immunocompetence and sperm performance demonstrated by Peters et al. (2004) in mallards; and Tomášek et al. (2016) use of pro-oxidant manipulations to reveal oxidative-status versus colour maintenance allocation-tradeoffs in captive zebra finches. Other comprehensive studies find clear links between oxidative stress, colour and sperm performance. In the zebra finch (*Taeniopygia guttata*), Tomášek et al. (2017) established that sperm performance of colourfully-beaked males was no different than drab males. However, dosing male zebra finch with the pro-oxidant diquat, the sperm performance of the most colourful males was drastically reduced in a remarkably similar fashion as the morph-specific relationship of superoxide and sperm performance in our current study (Figure 1 in Tomášek et al., 2017; our Figures 5a and 5b). Rojas Mora et al. (2016) found that the ratio of oxidised to reduced glutathione (an endogenous antioxidant that quenches hydrogen peroxide, a downstream product of superoxide dismutation by SOD) in the ejaculate of house sparrows (*Passer domesticus*) was negatively correlated with the size of a melanin-based breast badge, thus indicating a link between badge size and increased ejaculate antioxidants. Nevertheless, sperm swimming performance in the house sparrow was not directly related to any oxidant or antioxidant measurement in semen (Rojas Mora et al., 2016). In a follow up paper, experimental increases in oxidative stress in the same species of sparrow increased antioxidant levels in the ejaculate but diquat dosing still reduced sperm velocity with *the least* aggressive/dominant birds being the most strongly affected, probably due to socially mediated decreases in access to food resources—or condition dependence (Rojas Mora et al., 2017a). In yet another study, Rojas Mora et al 2017b, semen superoxide dismutase was positively correlated with blood SOD supporting our view that somatic oxidative status should relate to ejaculate oxidative status. Furthermore, males that changed social status to dominate roles tended to have lower quality ejaculates compared to those produced by males in the middle of the hierarchy (Figure 3, Rojas Mora et al., 2017b; unless the bird started at the lowest social status who then moved into higher ranks). Together Rojas Mora et al.’s comprehensive work reveals a complex relationship between oxidative status, ejaculate quality and social interactions in house sparrows, that we interpret broadly as pointing toward the extreme plasticity of ejaculate traits in response to social status and resource allocation trade-offs that are revealed under pro-oxidant manipulations, when the social system is perturbed, and when access to resources is limited by dominance related social interactions.

Initiating aggression is a critical determinant of winning male-male contests in painted dragons (Healey et al., 2007) and bibbed males are the more aggressive of the two bib-morphs (McDiarmid et al., 2017b), which may partially explain why males with bibs are less likely to lose paternity to neighboring rivals and males without bibs (Olsson et al., 2009a). Our results contrasts to some of the results found in the wild house sparrows described above; relative superoxide levels most strongly negatively associated with sperm performance of the more aggressive bibbed males, who also tended to have lower SOD in unmanipulated lizards. In our study, males were not allowed to interact (intense fighting occurs if males are housed together) which removes the costs of aggressive interactions (on both the aggressor and their target) and the lizards were fed to satiety every other day so socially mediated differential in resource limitation alone is not a likely explanation of our results. But if aggressiveness is correlated with underlying physiology, such as metabolic rate, costs may be incurred in addition to those due to direct interactions (Metcalfe et al., 1995). For example, aggressive male phenotypes of the Atlantic salmon (*Salmo salar*) have elevated metabolic rates which are beneficial in mating success, but may be deleterious during periods of low food abundance and even affect the growth rate and age-related declines in performance (Fleming, 1996; Jonsson et al., 1991; Metcalfe et al., 1995). Thus, the cost-benefit trade-off between morph-specific reproductive strategies may be linked to metabolic rate. It is possible, that such condition-dependent trade-offs underpin our results.

The correlations we found between ROS and sperm performance were in unmanipulated males, and although the relationship of body condition and SOD is only suggestive of condition-dependence. In the brown anole lizard (*Anolis sagrei*), experimental food-manipulations demonstrated body condition affected sperm morphology and numbers, which also translated to greater fertilization success for males with higher BCI (Kahrl and Cox, 2015). Manipulative studies such as those described above are necessary to fully illuminate the links we examine in this study.

Below we consider non-mutually exclusive hypotheses that may better explain our results while also suggesting further avenues of research. First, the temporal difference in these two condition dependent relationships with SOD, and the morph-specific differences in SOD may be related to the decrease in SOD and increase in mtSOx over the course of our sampling period. Second, morph-specific SOD turnover rates due to initial condition may have a lasting effect on SOD activity that may also explain the breakdown in the relationship between BCI and SOD at the time of mating. Regrettably, data on SOD turnover rates in animal taxa other than mammals (i.e., mice, rats and humans) are sparse and temperature dependent rates in ectotherms is in need of further research. Thermally mediated variation in SOD production and turnover rate was proposed as an explanation for the slower post activity decreases in mtSOx of male painted dragons acclimated to “hot” basking conditions compared with “cool” basking males (Ballen et al., 2012), but this was not tested directly. Nevertheless, we can construct rough estimates of turnover rates in these dragons because metabolic rate is linked to stoichiometry of protein and other macromolecular turnover across taxa (Hulbert and Else, 2000).

Some mammalian cellular proteins like histones (H3.1) and nuclear pore complex proteins (Nup205) persist nearly a year *in vivo* (Toyama et al., 2013), but most proteins have half-lives of only 45 minutes to 22.5 hours (Eden et al., 2011). Among proteins, superoxide dismutase is relatively long-lived (Valentine et al., 2005) with a half-life of 45-80 hours for SOD expressed in mice kidneys (Crisp et al., 2015; Hoffman et al., 1996). Nevertheless, the standard/basal metabolic rate of a 13g mouse (e.g., *Mus spp.*) is nearly 5.5 × higher than that of a 13 g painted dragon, (0.09ml/min*60 = 5.4ml/hr; Friesen et al., 2017a; mouse 29.78mlO2/hr; Withers, 1992, pg 95). Furthermore, non-specific protein synthesis rates in mice are ~4.8 × faster than in lizards (Sayegh and Lajtha, 1989; Warne et al., 2010) and isotopic nitrogen turnover rates yield similar values (Vander Zanden et al., 2015; Warne et al., 2010). Assuming that this inter-taxonomic factorial adjustment is reasonable, the degradation rate of SOD in a lizard is probably 4.8-5.5 × slower than in a mouse, which translates to a SOD half-life of 9-18 days in painted dragons. Given the upper end of this half-life range, BCI taken at capture could plausibly influence SOD activity 21-50 days later, which might explain the effect of early season condition on later SOD activity. As bibbed males have ~14% higher resting metabolic rates than non-bibbed males (Friesen et al., 2017a), they are likely to have had faster degradation of SOD, which could explain the disappearance of the positive relationship between SOD and BCI at mating in bibbed males, but not non-bibbed males.

Another, although more speculative, hypothesis is that different variants of SOD or mitochondrial genes may be linked to bib expression. For example, amyotrophic lateral sclerosis (ALS) is associated with oxidative damage of neuronal axons (Oeda et al., 2001), and mutations in SOD (>100 functional ALS mutations) are linked with 20-25% of ALS cases in humans (Valentine et al., 2005). In humans, small mutations in SOD genes results in drastic 10-30 fold increases in SOD levels in blood serum (Zelko et al., 2002). Thus, bibbed and non-bibbed males may have different genetic variants of SOD genes that affect oxidative status and is linked to condition. Nutrition and condition may also affect mitochondrial function and superoxide generation (Ballard and Youngson, 2015; Salin et al., 2018) and exogenous antioxidants could certainly play a role in reducing ROS levels (Monaghan et al., 2009). However, in painted dragons, supplementation with vitamin E (an antioxidant) does not increase longevity in the wild (Healey and Olsson, 2009b) and supplementation with dietary carotenoids does not reduce the effects of ROS effects on colour maintenance (Olsson et al., 2008a). Nevertheless, dietary antioxidants may affect sperm function in this species and this merits further research. We did not measure colour brightness or intensity in this study so we cannot directly assess the link between bib colour investment and sperm performance. McDiarmid et al. (2017b), however did not find a significant relationship between bib saturation and two measures of sperm function, motility (MOT) and velocity (VAP) which is consistent with the meta-analysis of Mautz et al. (2013) that found only weak positive, but non-significant correlations between colour traits and sperm kinematic performance. However, as Lüpold et al. (2014) demonstrate, the strength and evolution of pre- and postcopulatory trade-off depends on the degree of male monopolization. We might expect mate monopolization rates to be high for bibbed males based on advantages in both female preference and contest success (McDiarmid et al., 2017a), which seems to translate into reduced cuckoldry by neighbouring males in the wild (Olsson et al., 2009a). Nevertheless, if this trade-off is mediated by an allocation trade-off rather than a strict negative genetic correlation, as the relationship between SOD and body condition suggests it might, negative correlations between colouration and sperm performance might not be evident in the lab where resources were available ad libitum, and aggressive metabolically taxing territorial activity was absent.

Bibbed males have shorter telomeres than non-bibbed males (Rollings et al., 2017), and telomere shortening is linked with oxidative stress (Monaghan, 2010; Olsson et al., 2018; Olsson et al., 2017). In the wild, male painted dragons lose mass through the breeding season which ends in late December to mid-January; 90% of adults do not survive through winter ‘hibernation’ (beginning in late May) to the next breeding season (in September) (Healey and Olsson, 2009a; Olsson et al., 2007b; Olsson et al., 2009a). The increase in superoxide and decrease in SOD throughout the sampling period we describe here is consistent with senescence manifest in weight loss, DNA damage, telomere shortening, and colour fading across the breeding season in this species (Giraudeau et al., 2016; Olsson et al., 2012b; Rollings et al., 2017). We tested mtSOx in blood cells, but the mitochondrial proteins in sperm cells are the same as those in blood cells and should generate similar levels of mtSOx. Mitochondrial function and superoxide production is affected by the stress response (Manoli et al., 2007), and, if bib-morphs differ in their resilience to stress, this may explain different sensitivities of sperm to ROS. Spermatogenesis is ongoing throughout the breeding season in painted dragons, indicated by testicular enlargement persisting August through late December (Niejalke, 2006). Condition earlier in the season during spermatogenesis may have a strong influence on sperm damage and concomitant sensitivity to oxidative damage later in the season as our males were held without access to females prior to mating trials and sperm collection. This would constitute a trade-off between aggression typical of bibbed males and resilient sperm of non-bibbed males. Supplementation with a SOD mimetic, EUK 134, reduces age-related colour fading associated with oxidative stress and may also be used to experimentally tease apart the potential effects of body condition from SOD on sperm performance (Olsson et al., 2012b).

## Conclusions

We found a positive relationship between body condition and blood plasma SOD activity. SOD activity, which was lower in bibbed males, was negatively related to mitochondrial superoxide of erythrocytes. SOD was measured in the plasma which precludes a direct link between these variables. Furthermore, there was a negative, morph-specific indirect link between erythrocyte mitochondrial superoxide and sperm performance. Given the correlative nature of these results, we are unable to ascribe cause or disentangle the proximate mechanisms that underlay these the linkages. Nevertheless, these results add yet another morph-specific difference in physiology and behaviour to those we have previously documented in the Australian painted dragon lizards (Friesen et al., 2017a; Friesen et al., 2017b; Healey and Olsson, 2009a; Healey et al., 2007; McDiarmid et al., 2017a; Olsson et al., 2007a; Olsson et al., 2009a; Rollings et al., 2017). Future investigations that manipulate body condition and activity levels of bibbed and non-bibbed males are needed to determine whether sperm performance is condition-dependent in this species.

## Competing interests statement

We have no competing interests.

## Author contributions

All authors contributed materially and intellectually to this experiment and manuscript.

## Acknowledgements

Dr Danielle Johinke for tutorial on CASA and FACS. Dr Emily Uhrig, the Editor and three anonymous reviewers for useful and sagacious comments and edits. Dr Cissy Ballen for lizard husbandry and sperm collection. Dr Adele Haythornthwaite for administrative assistance. NSW National Parks service for permits and access to Yathong Nature Reserve, especially E. Snaith. National Science Foundation funding to CRF and Australian Research Council funding to SdG and MO. All research was conducted with the approval of the University of Sydney Animal Ethics Committee Project Number: 2013-2016/6050.

## Data, code and materials

Data will be deposited to http://datadryad.org/ upon acceptance of the manuscript.

## References

Aitken RJ, 2016. Oxidative stress and the etiology of male infertility. Journal of Assisted Reproduction and Genetics 33:1691–1692. doi: 10.1007/s10815-016-0791-4.

Aitken RJ, Clarkson JS, Fishel S, 1989. Generation of reactive oxygen species, lipid peroxidation, and human sperm function. Biol Reprod 41:183–197.

Aitken RJ, Graves JAM, 2002. Human spermatozoa: The future of sex. Nature 415:963–963.

Aitken RJ, Roman SD, 2008. Antioxidant systems and oxidative stress in the testes. Oxidative Medicine and Cellular Longevity 1:15–24.

Aitken RJ, Smith TB, Jobling MS, Baker MA, De Iuliis GN, 2014. Oxidative stress and male reproductive health. Asian Journal of Andrology 16:31.

Andreyev A, Kushnareva Y, Murphy A, Starkov A, 2015. Mitochondrial ROS metabolism: 10 years later. Biochemistry Biokhimiia 80:517.

Ballard JWilliam O, Youngson Neil A, 2015. Review: Can diet influence the selective advantage of mitochondrial DNA haplotypes? Bioscience Reports 35. doi: 10.1042/bsr20150232.

Ballen C, Healey M, Wilson M, Tobler M, Wapstra E, Olsson M, 2012. Net superoxide levels: Steeper increase with activity in cooler female and hotter male lizards. J Exp Biol 215:731–735.

Cogger HG, 2014. Reptiles and amphibians of Australia. Collingwood, VIC, Australia: CSIRO Publishing.

Costantini D, 2008. Oxidative stress in ecology and evolution: lessons from avian studies. Ecol Lett 11:1238–1251.

Costantini D, 2014. Oxidative stress and hormesis in evolutionary ecology and physiology : A marriage between mechanistic and evolutionary approaches. New York: Springer Science + Business Media.

Crisp MJ, Mawuenyega KG, Patterson BW, Reddy NC, Chott R, Self WK, Weihl CC, Jockel-Balsarotti J, Varadhachary AS, Bucelli RC, Yarasheski KE, Bateman RJ, Miller TM, 2015. In vivo kinetic approach reveals slow SOD1 turnover in the CNS. The Journal of Clinical Investigation 125:2772–2780. doi: 10.1172/JCI80705.

Dávila F, Aron S, 2017. Protein restriction affects sperm number but not sperm viability in male ants. J Insect Physiol 100:71–76.

De Graaf S, Evans G, Maxwell W, O’Brien J, 2006. In vitro characteristics of fresh and frozen–thawed ram spermatozoa after sex sorting and re-freezing. Reprod Fertil Dev 18:867–874.

delBarco-Trillo J, Roldan ERS, 2014. Effects of metabolic rate and sperm competition on the fatty-acid composition of mammalian sperm. J Evol Biol 27:55–62. doi: 10.1111/jeb.12275.

Dines JP, Mesnick SL, Ralls K, May-Collado L, Agnarsson I, Dean MD, 2015. A trade-off between precopulatory and postcopulatory trait investment in male cetaceans. Evolution 69:1560–1572. doi: 10.1111/evo.12676.

Dowling DK, Simmons LW, 2009. Reactive oxygen species as universal constraints in life-history evolution. Proc R Soc B 276:1737–1745.

Dunn JC, Halenar LB, Davies TG, Cristobal-Azkarate J, Reby D, Sykes D, Dengg S, Fitch WT, Knapp LA, 2015. Evolutionary trade-off between vocal tract and testes dimensions in howler monkeys. Curr Biol 25:2839–2844. doi: 10.1016/j.cub.2015.09.029.

Eden E, Geva-Zatorsky N, Issaeva I, Cohen A, Dekel E, Danon T, Cohen L, Mayo A, Alon U, 2011. Proteome half-life dynamics in living human cells. Science 331:764–768. doi: 10.1126/science.1199784.

Finkel T, Holbrook NJ, 2000. Oxidants, oxidative stress and the biology of ageing. Nature 408:239–247.

Firman RC, Simmons LW, 2010. Experimental evolution of sperm quality via postcopulatory sexual selection in house mice. Evolution 64:1245–1256.

Fitzpatrick JL, Almbro M, Gonzalez-Voyer A, Kolm N, Simmons LW, 2012. Male contest competition and the coevolution of weaponry and testes in pinnipeds. Evolution 66:3595–3604. doi: 10.1111/j.1558-5646.2012.01713.x.

Fitzpatrick JL, Lupold S, 2014. Sexual selection and the evolution of sperm quality. Mol Human Reprod 20:1180–1189. doi: 10.1093/molehr/gau067.

Fleming IA, 1996. Reproductive strategies of Atlantic salmon: ecology and evolution. Rev Fish Biol Fish 6:379–416.

Fridovich I, 1995. Superoxide radical and superoxide dismutases. Annu Rev Biochem 64:97–112.

Friesen CR, Johansson R, Olsson M, 2017a. Morph-specific metabolic rate and the timing of reproductive senescence in a color polymorphic dragon. J Exp Zool A Ecol Integr Physiol 327:433–443. doi: 10.1002/jez.2118.

Friesen CR, Shine R, Krohmer RW, Mason RT, 2013. Not just a chastity belt: the functional significance of mating plugs in garter snakes, revisited. Biol J Linn Soc 109:893–907. doi: 10.1111/bij.12089.

Friesen CR, Squire MK, Mason RT, 2014. Intrapopulational variation of ejaculate traits and sperm depletion in red-sided garter snakes. J Zool 292:192–201. doi: 10.1111/jzo.12092.

Friesen CR, Wilson MR, Rollings N, Sudyka J, Whittington CM, Giraudeau M, Olsson M, 2017b. Conditional handicaps in exuberant lizards: bright color in aggressive males is correlated with high levels of free radicals. Frontiers in Ecology and Evolution 5. doi: 10.3389/fevo.2017.00001.

Froman D, Feltmann A, 2000. Sperm mobility: Phenotype in roosters (Gallus domesticus) determined by concentration of motile sperm and straight line velocity. Biol Reprod 62:303–309.

Froman DP, Kirby JD, 2005. Sperm mobility: phenotype in roosters (Gallus domesticus) determined by mitochondrial function. Biol Reprod 72:562–567.

Garratt M, Bathgate R, de Graaf SP, Brooks RC, 2013. Copper-zinc superoxide dismutase deficiency impairs sperm motility and in vivo fertility. Reproduction 146:297–304.

Giraudeau M, Friesen CR, Sudyka J, Rollings N, Whittington CM, Wilson MR, Olsson M, 2016. Ageing and the cost of maintaining coloration in the Australian painted dragon. Biol Lett 12. doi: 10.1098/rsbl.2016.0077.

Gomendio M, Tourmente M, Roldan E, 2011. Why mammalian lineages respond differently to sexual selection: metabolic rate constrains the evolution of sperm size. Proc R Soc B 278:3135.

Healey M, Olsson M, 2009a. Too big for his boots: Are social costs keeping condition-dependent status signalling honest in an Australian lizard? Austral Ecol 34:636–640. doi: 10.1111/j.1442- 9993.2009.01968.x.

Healey M, Olsson M, 2009b. Vitamin E does not elevate survival in free-ranging lizards. Copeia 2009:339–341.

Healey M, Uller T, Olsson M, 2007. Seeing red: Morph-specific contest success and survival rates in a colour-polymorphic agamid lizard. Anim Behav 74:337–341.

Helfenstein F, Losdat S, Møller AP, Blount JD, Richner H, 2010. Sperm of colourful males are better protected against oxidative stress. Ecol Lett 13:213–222.

Hoffman EK, Wilcox HM, Scott RW, Siman R, 1996. Proteasome inhibition enhances the stability of mouse Cu Zn superoxide dismutase with mutations linked to familial amyotrophic lateral sclerosis. J Neurol Sci 139:15–20. doi: 10.1016/0022-510X(96)00031-7.

Hulbert A, Else PL, 2000. Mechanisms underlying the cost of living in animals. Annual Review of Physiology 62:207–235.

Huxley J, 1955. Morphism and evolution. Heredity 9:1–51.

Johinke D, de Graaf S, Bathgate R, 2014. Investigation of in vitro parameters and in vivo fertility of rabbit spermatozoa after chilled storage. Anim Reprod Sci 147:135–143.

Jonsson N, Hansen LP, Jonsson B, 1991. Variation in age, size and repeat spawning of adult Atlantic salmon in relation to river discharge. The Journal of Animal Ecology: 937–947.

Kahrl AF, Cox CL, Cox RM, 2016. Correlated evolution between targets of pre- and postcopulatory sexual selection across squamate reptiles. Ecol Evo 6:6452–6459.

Kahrl AF, Cox RM, 2015. Diet affects ejaculate traits in a lizard with condition-dependent fertilization success. Behav Ecol 26:1502–1511.

Ko EY, Sabanegh ES, Agarwal A, 2014. Male infertility testing: Reactive oxygen species and antioxidant capacity. Fertility and Sterility 102:1518–1527.

Koppers A, De Iuliis G, Finnie J, McLaughlin E, Aitken R, 2008. Significance of mitochondrial reactive oxygen species in the generation of oxidative stress in spermatozoa. J Clin Endocrin Met 93:3199.

Locatello L, Rasotto M, Evans J, Pilastro A, 2006. Colourful male guppies produce faster and more viable sperm. J Evol Biol 19:1595–1602.

Lüpold S, Tomkins JL, Simmons LW, Fitzpatrick JL, 2014. Female monopolization mediates the relationship between pre- and postcopulatory sexual traits. Nature Communications 5:1–8. doi: 10.1038/ncomms4184.

Lynch RE, Fridovich I, 1978. Permeation of the erythrocyte stroma by superoxide radical. J Biol Chem 253:4697–4699.

Manoli I, Alesci S, Blackman MR, Su YA, Rennert OM, Chrousos GP, 2007. Mitochondria as key components of the stress response. Trends in Endocrinology & Metabolism 18:190–198. doi: http://dx.doi.org/10.1016/j.tem.2007.04.004.

Mattson KJ, Vries AD, McGuire SM, Krebs J, Louis EE, Loskutoff NM, 2007. Successful artificial insemination in the corn snake, Elaphe gutatta, using fresh and cooled semen. Zoo Biol 26:363–369.

Mautz BS, Møller AP, Jennions MD, 2013. Do male secondary sexual characters signal ejaculate quality? A meta-analysis. Biol Rev 88:669–682.

McDiarmid CS, Friesen CR, Ballen C, Olsson M, 2017a. Sexual coloration and sperm performance in the Australian painted dragon lizard, Ctenophorus pictus. J Evol Biol 30:1303–1312. doi: 10.1111/jeb.13092.

McDiarmid CS, Friesen CR, Ballen C, Olsson M, 2017b. Sexual coloration and sperm performance in the Australian painted dragon lizard, Ctenophorus pictus. J Evol Biol:n/a-n/a. doi: 10.1111/jeb.13092.

Melville J, Schulte II JA, 2001. Correlates of active body temperatures and microhabitat occupation in nine species of central Australian agamid lizards. Austral Ecol 26:660–669. doi: 10.1046/j.1442-9993.2001.01152.x.

Metcalfe NB, Alonso-Alvarez C, 2010. Oxidative stress as a life-history constraint: The role of reactive oxygen species in shaping phenotypes from conception to death. Funct Ecol 24:984–996.

Metcalfe NB, Taylor AC, Thorpe JE, 1995. Metabolic rate, social status and life-history strategies in Atlantic salmon. Anim Behav 49:431–436. doi: http://dx.doi.org/10.1006/anbe.1995.0056.

Mitre R, Cheminade C, Allaume P, Legrand P, Legrand AB, 2004. Oral intake of shark liver oil modifies lipid composition and improves motility and velocity of boar sperm. Theriogenology 62:1557–1566.

Møller AP, 1988. Ejaculate quality, testes size and sperm competition in primates. J Hum Evol 17:479– 488.

Monaghan P, 2010. Telomeres and life histories: The long and the short of it. Ann N Y Acad Sci 1206:130–142. doi: 10.1111/j.1749-6632.2010.05705.x.

Monaghan P, Costantini D, 2014. Free Radicals–An Evolutionary Perspective. Systems Biology of Free Radicals and Antioxidants: Springer. p. 39–64.

Monaghan P, Metcalfe NB, Torres R, 2009. Oxidative stress as a mediator of life history trade-offs: Mechanisms, measurements and interpretation. Ecol Lett 12:75–92. doi: DOI 10.1111/j.1461-0248.2008.01258.x.

Murphy MP, 2009. How mitochondria produce reactive oxygen species. Biochemical Journal 417:1–13.

Niejalke DP, 2006. Reproduction by a small agamid lizard, Ctenophorus pictus, during contrasting seasons. Herpetologica 62:409–420.

Noguera JC, Dean R, Isaksson C, Velando A, Pizzari T, 2012. Age-specific oxidative status and the expression of pre-and postcopulatory sexually selected traits in male red junglefowl, Gallus gallus. Ecol Evo.

Oeda T, Shimohama S, Kitagawa N, Kohno R, Imura T, Shibasaki H, Ishii N, 2001. Oxidative stress causes abnormal accumulation of familial amyotrophic lateral sclerosis-related mutant SOD1 in transgenic Caenorhabditis elegans. Hum Mol Genet 10:2013–2023.

Olsson M, 2001. ’Voyeurism’ prolongs copulation in the dragon lizard Ctenophorus fordi. Behav Ecol Sociobiol 50:378–381.

Olsson M, Friesen CR, Rollings N, Sudyka J, Lindsay W, Whittingtion CM, Wilson M, 2018. Long term effects of superoxide and DNA repair on lizard telomeres. Mol Ecol 0. doi: doi:10.1111/mec.14913.

Olsson M, Healey M, Astheimer L, 2007a. Afternoon T: Testosterone level is higher in red than yellow male polychromatic lizards. Physiol Behav 91:531–534.

Olsson M, Healey M, Perrin C, Wilson M, Tobler M, 2012a. Sex-specific SOD levels and DNA damage in painted dragon lizards (Ctenophorus pictus). Oecologia 170:917–924. doi: 10.1007/s00442-012-2383-z.

Olsson M, Healey M, Wapstra E, Schwartz T, Lebas N, Uller T, 2007b. Mating system variation and morph fluctuations in a polymorphic lizard. Mol Ecol 16:5307–5315.

Olsson M, Healey M, Wapstra E, Uller T, 2009a. Testing the quality of a carrier: a field experiment on lizard signalers. Evolution International Journal of Organic Evolution 63:695.

Olsson M, Schwartz T, Uller T, Healey M, 2009b. Effects of sperm storage and male colour on probability of paternity in a polychromatic lizard. Anim Behav 77:419–424.

Olsson M, Tobler M, Healey M, Perrin C, Wilson M, 2012b. A significant component of ageing (DNA damage) is reflected in fading breeding colors: An experimental test using innate antioxidant memetics in painted dragon lizards. Evolution 66:2475–2483. doi: 10.1111/j.1558-5646.2012.01617.x.

Olsson M, Wapstra E, Friesen CR, 2017. Evolutionary Ecology of Telomeres: a Review. Ann N Y Acad Sci In press.

Olsson M, Wilson M, Isaksson C, Uller T, 2009c. Polymorphic ROS scavenging revealed by CCCP in a lizard. Naturwissenschaften 96:845–849.

Olsson M, Wilson M, Isaksson C, Uller T, Mott B, 2008a. Carotenoid intake does not mediate a relationship between reactive oxygen species and bright colouration: Experimental test in a lizard. J Exp Biol 211:1257–1261.

Olsson M, Wilson M, Uller T, Mott B, Isaksson C, Healey M, Wanger T, 2008b. Free radicals run in lizard families. Biol Lett 4:186–188.

Parapanov R, Nusslé S, Hausser J, Vogel P, 2008. Relationships of basal metabolic rate, relative testis size and cycle length of spermatogenesis in shrews (Mammalia, Soricidae). Reprod Fertil Dev 20:431.

Parker GA, 1970. Sperm competition and its evolutionary consequences in insects. Biol Rev Camb Philos Soc 45:525-&. doi: 10.1111/j.1469-185X.1970.tb01176.x.

Parker GA, Lessells CM, Simmons LW, 2013. Sperm competition games: a general model for precopulatory male-male competition. Evolution 67:95–109. doi: 10.1111/j.1558-5646.2012.01741.x.

Parker GA, Pizzari T, 2010. Sperm competition and ejaculate economics. Biological Reviews 85:897–934. doi: 10.1111/j.1469-185X.2010.00140.x.

Peters A, Denk A, Delhey K, Kempenaers B, 2004. Carotenoid-based bill colour as an indicator of immunocompetence and sperm performance in male mallards. J Evol Biol 17:1111–1120.

Pitcher T, Rodd F, Rowe L, 2007. Sexual colouration and sperm traits in guppies. Journal of Fish Biology 70:165–177.

Pryke SR, Astheimer LB, Buttemer WA, Griffith SC, 2007. Frequency-dependent physiological trade-offs between competing colour morphs. Biol Lett 3:494–497. doi: 10.1098/rsbl.2007.0213.

Pryke SR, Griffith SC, 2006. Red dominates black: agonistic signalling among head morphs in the colour polymorphic Gouldian finch. Proc R Soc B 273:949–957. doi: 10.1098/rspb.2005.3362.

Quintero-Moreno A, Miró J, Teresa Rigau A, Rodríguez-Gil JE, 2003. Identification of sperm subpopulations with specific motility characteristics in stallion ejaculates. Theriogenology 59:1973–1990. doi: http://dx.doi.org/10.1016/S0093-691X(02)01297-9.

Rakitin A, Ferguson MM, Trippel EA, 1999. Sperm competition and fertilization success in Atlantic cod (Gadus morhua): effect of sire size and condition factor on gamete quality. Can J Fish Aquat Sci 56:2315–2323.

Ramm SA, Schärer L, 2014. The evolutionary ecology of testicular function: Size isn’t everything. Biol Rev 89:874–888.

Ribou A-C, Reinhardt K, 2012. Reduced metabolic rate and oxygen radicals production in stored insect sperm. Proc R Soc B 279:2196–2203. doi: 10.1098/rspb.2011.2422.

Rickard JP, Pini T, Soleilhavoup C, Cognie J, Bathgate R, Lynch GW, Evans G, Maxwell W, Druart X, De Graaf S, 2014. Seminal plasma aids the survival and cervical transit of epididymal ram spermatozoa. Reproduction 148:469–478.

Rojas Mora A, Firth A, Blareau S, Vallat A, Helfenstein F, 2017a. Oxidative stress affects sperm performance and ejaculate redox status in subordinate house sparrows. J Exp Biol:jeb. 154799.

Rojas Mora A, Meniri M, Glauser G, Vallat A, Helfenstein F, 2016. Badge size reflects sperm oxidative status within social groups in the House Sparrow Passer domesticus. Frontiers in Ecology and Evolution 4:67.

Rojas Mora A, Meniri M, Gning O, Glauser G, Vallat A, Helfenstein F, 2017b. Antioxidant allocation modulates sperm quality across changing social environments. PLoS ONE 12:e0176385.

Rollings N, Friesen CR, Sudyka J, Whittington C, Giraudeau M, Wilson M, Olsson M, 2017. Telomere dynamics in a lizard with morph-specific reproductive investment and self-maintenance. Ecol Evo 7:5163–5169. doi: 10.1002/ece3.2712.

Rowe M, Swaddle JP, Pruett-Jones S, Webster MS, 2010. Plumage coloration, ejaculate quality and reproductive phenotype in the red-backed fairy-wren. Anim Behav 79:1239–1246. doi: 10.1016/j.anbehav.2010.02.020.

Rutherford A, 2011. ANOVA and ANCOVA: a GLM approach: John Wiley & Sons.

Salin K, Villasevil EM, Anderson GJ, Auer SK, Selman C, Hartley RC, Mullen W, Chinopoulos C, Metcalfe NB, 2018. Decreased mitochondrial metabolic requirements in fasting animals carry an oxidative cost. Funct Ecol 32:2149–2157. doi:doi:10.1111/1365-2435.13125.

Sanocka D, Kurpisz M, 2004. Reactive oxygen species and sperm cells. Reproductive Biology and Endocrinology 2:1–7.

Sayegh JF, Lajtha A, 1989. In vivo rates of protein synthesis in brain, muscle, and liver of five vertebrate species. Neurochem Res 14:1165–1168. doi: 10.1007/bf00965625.

Simmons LW, Emlen DJ, 2006. Evolutionary trade-off between weapons and testes. Proc Natl Acad Sci USA 103:16346–16351.

Simmons LW, Firman RC, Rhodes G, Peters M, 2004. Human sperm competition: testis size, sperm production and rates of extrapair copulations. Anim Behav 68:297–302.

Simmons LW, Fitzpatrick JL, 2012. Sperm wars and the evolution of male fertility. Reproduction 144:519–534. doi: 10.1530/rep-12-0285.

Simon L, Lewis SEM, 2011. Sperm DNA damage or progressive motility: which one is the better predictor of fertilization in vitro? Systems Biology in Reproductive Medicine 57:133–138. doi: 10.3109/19396368.2011.553984.

Sinervo B, Lively CM, 1996. The rock-paper-scissors game and the evolution of alternative male strategies. Nature 380:240–243. doi: 10.1038/380240a0.

Svensson PA, Wong B, 2011. Carotenoid-based signals in behavioural ecology: A review. Behaviour 148:131–189.

Tazzyman SJ, Pizzari T, Seymour RM, Pomiankowski A, 2009. The evolution of continuous variation in ejaculate expenditure strategy. Am Nat 174:E71–E82.

Tomášek O, Albrechtová J, Němcová M, Opatová P, Albrecht T, Trade-off between carotenoid-based sexual ornamentation and sperm resistance to oxidative challenge. Proc R Soc B 2017. The Royal Society. p. 20162444.

Tomášek O, Gabrielová B, Kačer P, Maršík P, Svobodová J, Syslová K, Vinkler M, Albrecht T, 2016. Opposing effects of oxidative challenge and carotenoids on antioxidant status and condition-dependent sexual signalling. Scientific reports 6.

Tourmente M, Gomendio M, Roldan ER, 2011. Mass-specific metabolic rate and sperm competition determine sperm size in marsupial mammals. PLoS ONE 6:e21244.

Toyama Brandon H, Savas Jeffrey N, Park Sung K, Harris Michael S, Ingolia Nicholas T, Yates John R, Hetzer Martin W, 2013. Identification of long-lived proteins reveals exceptional stability of essential cellular structures. Cell 154:971–982. doi: http://dx.doi.org/10.1016/j.cell.2013.07.037.

Tremellen K, 2008. Oxidative stress and male infertility--a clinical perspective. Human Reproduction Update 14:243.

Tuni C, Han CS, Dingemanse NJ, 2018. Multiple biological mechanisms result in correlations between pre-and post-mating traits that differ among versus within individuals and genotypes. Proc R Soc B 285:20180951.

Turrens JF, 2003. Mitochondrial formation of reactive oxygen species. The Journal of Physiology 552:335–344.

Valentine JS, Doucette PA, Zittin Potter S, 2005. Copper-zinc superoxide dismutase and amyotrophic lateral sclerosis. Annu Rev Biochem 74:563–593.

van der Horst G, Maree L, 2014. Sperm form and function in the absence of sperm competition. Mol Reprod Dev 81:204–216.

Vander Zanden MJ, Clayton MK, Moody EK, Solomon CT, Weidel BC, 2015. Stable isotope turnover and half-life in animal tissues: A literature synthesis. PLoS ONE 10:e0116182.

von Schantz T, Bensch S, Grahn M, Hasselquist D, Wittzell H, 1999. Good genes, oxidative stress and condition–dependent sexual signals. Proc R Soc B 266:1–12.

Warne RW, Gilman CA, Wolf BO, 2010. Tissue-carbon incorporation rates in lizards: implications for ecological studies using stable isotopes in terrestrial ectotherms. Physiol Biochem Zool 83:608–617.

Wellenreuther M, Svensson EI, Hansson B, 2014. Sexual selection and genetic colour polymorphisms in animals. Mol Ecol 23:5398–5414.

Withers P, 1992. Comparative Animal Physiology. Philadephia: Saunders College Publishing.

Zelko IN, Mariani TJ, Folz RJ, 2002. Superoxide dismutase multigene family: a comparison of the CuZn-SOD (SOD1), Mn-SOD (SOD2), and EC-SOD (SOD3) gene structures, evolution, and expression. Free Radical Biol Med 33:337–349.

